# Flexiplex: A versatile demultiplexer and search tool for omics data

**DOI:** 10.1101/2023.08.21.554084

**Authors:** Oliver Cheng, Min Hao Ling, Changqing Wang, Shuyi Wu, Matthew E. Ritchie, Jonathan Göke, Noorul Amin, Nadia M. Davidson

## Abstract

The process of analyzing high throughput sequencing data often requires the identification and extraction of specific target sequences. This could include tasks such as identifying cellular barcodes and UMIs in single cell data, and specific genetic variants for genotyping. However, existing tools which perform these functions are often task-specific, such as only demultiplexing barcodes for a dedicated type of experiment, or are not tolerant to noise in the sequencing data. To overcome these limitations, we developed Flexiplex, a versatile and fast sequence searching and demultiplexing tool for omics data, which is based on the Levenshtein distance and thus allows imperfect matches. We demonstrate Flexiplex’s application on three use cases, identifying cell line specific sequences in Illumina short-read single cell data, and discovering and demultiplexing cellular barcodes from noisy long-read single cell RNA-seq data. We show that Flexiplex achieves an excellent balance of accuracy and computational efficiency compared to leading task-specific tools. Flexiplex is available at https://davidsongroup.github.io/flexiplex/.

## Introduction

High throughput sequencing, from both short-read and long-read sequencing technologies, enable the genome, transcriptome and other ‘omics to be profiled by reading up to billions of nucleotides. Matching and extracting sequences from ‘omics data is a common analysis task. For example, searching for the presence of genetic variants or motifs, or identifying and error correcting barcodes. Highly efficient string search command line tools such as grep and agrep (Wu and Manber), which find exact and approximate matches respectively, have been used for this purpose in the past (Panagopoulos *et al*., 2014). However, they are designed for a small set of search patterns, and do not output the data in a format compatible with downstream analysis tools, thereby limiting their use for demultiplexing. Demultiplexing is becoming increasingly routine in ‘omics data analysis. For instance, indexes, including barcodes and Unified Molecular Identifiers (UMIs), are added for a range of purposes such as pooling samples to save sequencing cost, measuring clonal expansion, labeling reads from individual cells, or individual molecules (Smith *et al*., 2010; Bramlett *et al*., 2020; Merino *et al*., 2019; Philpott *et al*., 2021). To address this, demultiplexing tools have been built such as ultraplex (Wilkins *et al*., 2021) for short-read data, and scTagger (Ebrahimi *et al*., 2022) and BLAZE (You et al., 2023) for noisy long-read data. Some demultiplexing tools have also been built into multi-purpose pipelines such as FLAMES (Tian *et al*., 2021), wf-single-cell (https://github.com/epi2me-labs/wf-single-cell) and SiCeLoRe (Lebrigand *et al*., 2020). However these tools are often tailored to a specific experiment type, and may not work on a wider range of customized barcodes and sequencing data, specifically, when the structure of the barcode, flanking sequences, and their locations differ from the usual specifications. For this reason, experiment-agnostic tools have begun to be developed (Sullivan and Pachter, 2023).

Long-read demultiplexing is a particular challenge, as the reads often contain a high number of errors, including insertions and deletions (Dohm *et al*., 2020). Moreover, a small percent of reads may be chimeric (White *et al*., 2017), containing multiple barcodes and requiring read to be split. Although there are some tools which can simultaneously split chimeric reads and demultiplex, such as poreChop (Wick, 2017), these may not work across a broad range of datasets or when there are thousands of barcodes. Finally, much of the available software is complex to setup and install and their computational requirements in terms of time and memory can limit their practical use when processing the volumes of data now being generated.

Here we introduce Flexiplex, which, given a set of reads as either FASTQ or FASTA, will demultiplex and/or identify a sequence of interest, reporting matching reads and read-barcode assignment. Flexiplex works in two modes: a) when one or more sequences of interest are known, such as barcodes and b) discovery mode – when only the sequence which flanks the region of interest is known. Flexiplex is lightweight, multithreaded, fast, and easy to install and run. The source code and pre-built binary executables for Linux and MacOS are provided at https://davidsongroup.github.io/flexiplex/.

### Flexiplex’s Approach

Flexiplex assumes a read structure where a barcode and UMI are flanked by other known sequences (Figure 1A). To identify the barcode, Flexiplex first uses the edlib C++ library (Šošić and Šikić, 2017) to search for the left and/or right flanking sequence. Default sequences are 22bp (base pairs) from the 10x Genomics Chemistry v3 primer and 9bp of poly T sequence. The intermediate sequence which contains the barcode and UMI, is initially left as a wildcard (default length of 28bp). For the best match to the flanking sequence within a specified edit distance (default edit distance of 8), the intermediate sequence (barcode and UMI) +/-5 bp either side is extracted. Next, we align the extracted sequence against a user-provided list of known barcodes and calculate the Levenshtein distance (Berger *et al*., 2021). For speed, this step is implemented in an efficient dynamic programming algorithm within Flexiplex, rather than edlib. The best matching barcode equal or less than a specified distance (default of two) is reported, but only if no other barcode has equal lowest distance. To identify and split chimeric reads, Flexiplex will repeat the flank and barcode search with the previously found barcode and flanking region masked out. This is repeated until no new barcodes are found in the read. Subsequently, the reverse complement of the unmasked read is searched using the same algorithm.

**Figure 1:**
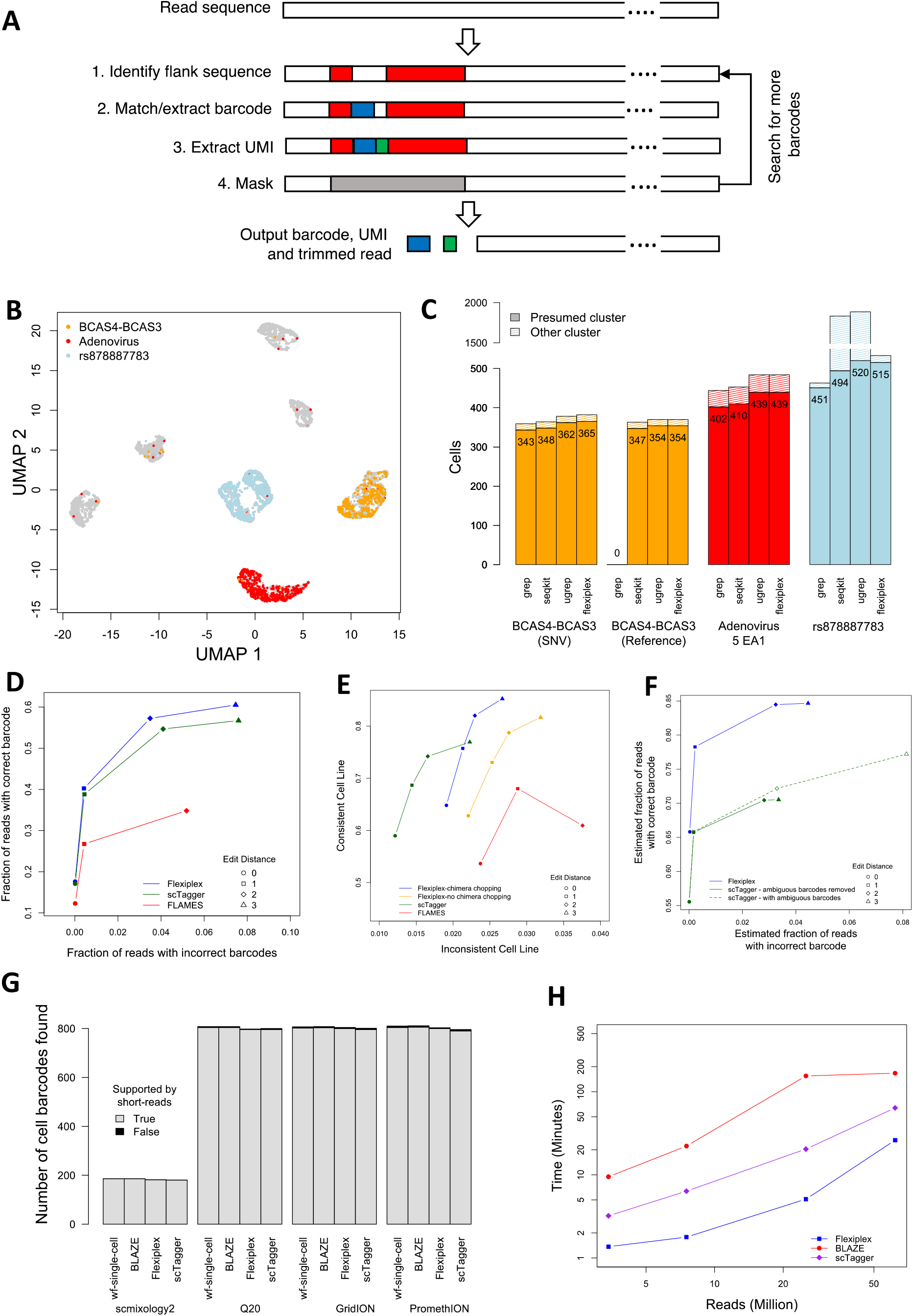
(A) The demultiplexing approach used by Flexiplex. The right and left flank are first searched for within a read. The barcode and UMI regions are then extracted from the intermediate sequence, with barcode error correction if known barcodes are provided. (B) UMAP of the short-read single cell dataset of 7 pooled cell lines. Cells positive for BCAS4-BCAS3 (orange), Adenovirus 5 EA1 (red) and rs878887783 (blue) are indicated. (C) The number of cells identified with grep, seqkit grep, ugrep and Flexiplex that express sequence from BCAS4-BCAS3 (SNP - using an MCF-7 specific variant or Reference - using the reference allele), Adenovirus 5 EA1 and rs878887783 in a short-read single cell dataset of 7 pooled cells lines. Cells which cluster away from the presumed cluster (hatched) are likely to be false positives, whereas those falling within the presumed cluster are true positives (values on bars) (D) The accuracy of barcode demultiplexing on a simulated set of 5 million single cell RNA-seq long-reads for Flexiplex, scTagger and FLAMES, varying the maximum allowed edit distance to known barcodes between 0 to 3. (E) Assessment of cellular barcode demultiplexing on a real dataset of 248 cells sequenced with ONT for Flexiplex (with and without chimeric read splitting), scTagger and FLAMES, varying the maximum allowed edit distance to known barcodes between 0 to 3. Correct barcodes will result in a higher level of consistent cell line annotation. (F) Performance of Flexiplex and scTagger on a large dataset of 61 million reads, where decoy barcodes were used to assess demultiplexing accuracy. As scTagger reports multiple barcodes of equi-distance for each read, we assessed its performance by either removing reads with ambiguous reads, or counting any true barcode as a true positive. (G) The number of barcodes recovered across four datasets when no known barcode list was provided. As scTagger does not adjust the produced barcodes to remove empty droplets like the other methods, we used a script provided with Flexiplex, flexiplex-filter, to automatically refine the barcodes based on the end of the inflection point of the read-barcode frequency distribution. (H) The run-time (log scale, four threads) of stand-alone tools for barcode discovery, Flexiplex, BLAZE and scTagger, as a function of the number of reads processed from the four datasets used for barcode discovery evaluation. See text and Supplementary Material for further details.

The flank and barcode sequences provided to Flexiplex are customisable, as is the length of the barcode and UMI, and their order. Flexiplex will output identified barcodes, UMIs, and trimmed reads by default. If the barcode, UMI and right flank are left unspecified, Flexiplex can be used as a general searching tool and will identify any read containing the left flanking sequence within a user defined distance. As it can read and write to standard IO, the output of Flexiplex can be piped to another instance of Flexiplex for complex search and demultiplexing applications, such as extracting cell barcodes from all reads which contain a genetic variant of interest.

Flexiplex can also be used to discover barcodes directly from the data. If no barcode list is provided, the sequence following the left flank is assumed as the barcode, and barcodes are reported, along with their frequencies. Flexiplex is provided with a Python script, flexiplex-filter, which generates a filtered barcode list based on knee plot frequencies and, optionally, a user-provided whitelist. Flexiplex-filter works by first using a rolling window to approximate the derivative of the knee plot curve for each barcode. The most negative derivative is then assumed as the inflection point, which becomes the cutoff point. A plot of the frequency distribution and inflection point can be generated for manual inspection and the search range customized if needed (Supplementary Figure 1). The filtered barcodes can then be passed back into Flexiplex for error tolerant barcode demultiplexing of each read.

To illustrate the performance of Flexiplex, we considered several typical use cases; searching for specific sequences of interest in low error short-read data and discovering and demultiplexing single cell RNA sequencing barcodes from noisy long-read data. In all instances Flexiplex demonstrated an excellent balance between computational efficiency and accuracy. However these use cases are not exhaustive. Flexiplex is versatile, with a wide range of applications beyond the examples presented here, such as clonal tagging of single cells using barcoding approaches (Putri *et al*., 2023) and sample demultiplexing in bulk data.

## Use Cases

### Fast and accurate sequence search

We first show Flexiplex is a reliable and general sequence searching tool using an Illumina short-read single cell RNA-Seq dataset from Chen et al. (Chen *et al*., 2021). This data was generated from a mixture of 7 cell lines including MCF-7 which is known to harbor the highly expressed fusion gene BCAS4-BCAS3 (Edgren et al., 2011; Davidson *et al*., 2015). The pool also included the HEK293T cell line which expresses E1A, an adenovirus 5 gene and the T47D cell line which contains a Single Nucleotide Variant (SNV), rs878887783, in the mitochondrial gene, MT-ND1. Here we demonstrate how reads of interest can be extracted using a query sequence and then processed for downstream analysis; in this use case, identifying reads from BCAS4-BCAS3, Adenovirus E1A and rs878887783, and using them to assign cells to their cell line of origin.

Using Flexiplex, we searched the reads for 34bp from each of Adenovirus E1A and the dominant breakpoint of BCAS4-BCAS3, and 54bp centered on rs878887783 within an edit distance of two. As BCAS4-BCAS3 has an MCF-7 specific SNV 13bp from the breakpoint, we used two query sequences, one with the SNV and one with the reference allele. We compared these results against three alternative methods: grep, which looks for an exact match; and ugrep (https://github.com/Genivia/ugrep) and seqkit grep (Shen *et al*., 2016), which both allow mismatches. Flexiplex processed approximately 200 million reads in 24 minutes using a single thread per search (linux high-performance computer). The computational speed was slower than grep (which took 1-2 minutes), but significantly faster than the other mismatch tolerant methods, seqkit grep (∼190 minutes) and ugrep (∼40 minutes) (Supplementary Figure 2).

To translate identified reads into a list of cells expressing the novel sequence, we extracted cellular barcodes (first 16bp) from the pair of matched reads. The cells identified fell into clusters based on gene expression as expected (Figure 1B), allowing us to assign clusters to cell lines and assess the performance of each tool. Flexiplex identified the highest or second highest number of cells expressing BCAS4-BCAS3 (365 of 956 cells), Adenovirus E1A (439 of 869 cells) and rs878887783 (515 of 987 cells) in the correct cluster (Figure 1C). The importance of allowing mismatches was exemplified by BCAS4-BCAS3. When using the reference allele in our query sequence rather than the MCF-7 specific SNP, grep reported no matching reads, whereas Flexiplex identified 97% of those found using the MCF-7 variant (Figure 1C).

False positives for BCAS4-BCAS3 and Adenovirus E1A, assessed by cells expressing the novel genes from other cell line clusters, were at a similar rate for all tools. We hypothesize that the false positives are unlikely to be a result of poor sequence matching, but rather, a result of sequencing errors in the cellular barcodes, ambient RNA (Young and Behjati, 2020) or barcode switching (Griffiths *et al*., 2018). To confirm this, we searched for the same variants in bulk RNA sequencing from three of the same cell lines and found no false positives for BCAS4-BCAS3 or Adenovirus E1A (Supplementary Table 1). For rs878887783, however, a large number of false positives were seen for ugrep and seqkit in both single cell (Figure 1C, Supplementary Figure 3) and bulk (Supplementary Table 1) data. These can be attributed to detection of the reference allele in other cell lines. As Flexiplex allows different error tolerance for different regions of the search sequence, it could require a perfect match for the 5bp at the SNV while allowing up to 2 mismatches in the flanking region. Taken together, these results demonstrate that error tolerance improves sensitivity without compromising precision, even for low error Illumina data.

### Demultiplexing cellular barcodes from noisy long-read data

Barcode demultiplexing is an application of sequence searching, where the number of query sequences (barcodes) can be large and the best match should be reported. We validated Flexiplex’s demultiplexing performance on noisy long-read Oxford Nanopore Technology (ONT) single cell RNA sequencing. We compared Flexiplex’s performance with two specialized tools for long-read single cell demultiplexing, scTagger and FLAMES, on simulated data generated by Ebrahimi et al (Ebrahimi *et al*., 2022). The simulation consisted of 5 million reads generated for 5 thousand cells. True barcode sequences were provided to each tool to benchmark them under a best-case scenario, where a perfect list of known barcodes is provided. Flexiplex reported the correct barcodes for more reads than any other methods across all maximum edit distances tested (0-3) and had the lowest false discovery rate (Figure 1D).

Next, we verified these results on a real dataset, scmixology 2 from Tian et al. (Tian *et al*., 2021). Scmixology 2 is a pool of five cell lines which underwent FLT-Seq (Jabbari and Tian), where cells were processed with the 10x Genomics 3’ v3 protocol followed by ONT long-read cDNA sequencing in addition to Illumina short-read sequencing. Approximately 25 million ONT reads were generated. 248 cells were identified in the matched short-read sequencing and their barcodes passed to the long-read demultiplexing tools.

To test the cellular barcode demultiplexing accuracy, we used two orthogonal methods to determine a read’s cell line of origin and assess their concordance. In the first method, cells had previously been annotated to cell lines using SNP-based clustering from matched short-reads (Tian *et al*., 2021). Reads could therefore be assigned to cell lines by correctly demultiplexing their cellular barcodes and looking up their associated cell line from the annotation. For a subset of reads, we could also determine the cell line using a second method, using cell line specific Single Nucleotide Polymorphisms (SNPs) present in the reads (16 thousand reads). For each of these SNP-typed reads, we assessed whether the predicted cell line from each approach matched (See Supplementary Methods for details), with the assumption that inconsistencies were predominantly caused by incorrect cellular barcodes.

We found that Flexiplex reported the highest number of reads where the cell lines were concordant, whilst less than 5% of reads had discordant cell lines (Figure 1E). Importantly, we found that 3% of the reads in the dataset were chimeric, and Flexiplex’s ability to split these improved the demultiplexing performance further. Flexiplex took 2.6 hours to process 25 million long reads (default settings, single thread, linux high-performance computer), which was slower than FLAMES (1.1 hours), but faster than scTagger (2.9 hours). Both Flexiplex and FLAMES consumed less than 1 GB of RAM, compared to 41 GB for scTagger (Supplementary Figure 4).

Finally, we verified the performance of Flexiplex on a large PromethION dataset from You et al. (You *et al*., 2023) of over one thousand human induced pluripotent stem cells (hiPSC) sequenced to a depth of 61 million reads. To emulate a typical 10x Genomics experiment, we passed each tool approximately 11 thousand cellular barcodes, where 1022 were true barcodes obtained from short-read data and the remaining 10,000 were decoys -randomly sampled from 10x Genomics’ list of possible cellular barcodes for v3. We then assessed the rates that true and decoy barcodes were reported (see Supplementary Methods) and found that Flexiplex was able to demultiplex the highest number of reads correctly, consistent with other analyses (Figure 1F). Flexiplex completed in 7-14 hours using 16 threads. scTagger was considerably faster, taking just 1.5-5 hours, but required over 100 GB of RAM, compared to < 1 GB for Flexiplex. FLAMES was unable to complete in a reasonable time due to being single threaded (< 30% complete after 48 hours), so was excluded from the comparison.

### Discovering cellular barcode from noisy long-read data

Next we assessed Flexiplex’s performance at barcode discovery, a scenario where the barcodes are unknown, and a list of valid barcodes need to be generated prior to demultiplexing with tools such as those in the previous use case. This is one of the first analysis steps required for long-read single-cell sequencing without matched short-reads. We compared Flexiplex’s barcode discovery against the purpose built tools scTagger, BLAZE, and ONT’s wf-single-cell on the scmixology 2 dataset, as well as three technical replicates (GridION Q20, GridION and PromethION) of human induced pluripotent stem cells (hiPSC) sequenced with varying depths and error rates by You et al. (You *et al*., 2023). The associated barcodes derived from short-read data were used as truth, but not provided to the tools. As scTagger does not estimate the number of cells using a knee plot method, we applied flexiplex-filter to the barcode frequencies from scTagger, to obtain the short-list for comparison. All tools were found to have similar sensitivity and specificity (Figure 1G). However, we found that Flexiplex (4 threads, other settings default, Linux high-performance computer) completed barcode discovery in approximately 5-16% of the time of BLAZE and 25-42% of the time of scTagger (Figure 1H). Relative performance was similar for other thread counts (Supplementary Table 2). While wf-single-cell is able to generate a barcode list, it is a complete Netflow pipeline for single-cell data preprocessing, and will therefore perform tasks unrelated to barcode discovery. Hence computational performance will exceed other tools and cannot be compared directly. For example, wf-single-cell (16 threads, up to 20 concurrent jobs) required memory and runtime an order of magnitude higher than Flexiplex (Supplementary Table 2).

## Discussion

Here we present Flexiplex, a generalized tool for sequence searching and demultiplexing. To achieve computational efficiency on large ‘omics data, Flexiplex uses a combination of two Levenshtein distance alignment algorithms. Hence Flexiplex is tolerant to substitutions, insertions and deletions which can be present in the data due to sequencing errors and SNPs. Flexiplex is highly customisable, including all search sequences and lengths, and whether to split chimeric reads, making it a versatile tool for many applications.

Using barcode discovery and demultiplexing of single cell long-read RNA-Seq, and sequence searching in single cell short-read RNA-Seq as use cases, we demonstrate that Flexiplex achieves accuracy comparable or exceeding popular task-specific tools, with good run-time and memory usage. However, Flexiplex’s applications are broader - beyond single cell and RNA sequencing. For example, demultiplexing pooled samples generated by bulk long read experiments, or custom clonal barcodes. Because Flexiplex can read and write to standard IO, instances of itself can be chained together on the command line for more complex tasks, for example, to select reads with a mutation of interest and then demultiplex their cellular barcodes, or to search for phased variants or splicing. We designed Flexiplex to be a simple, user-friendly and fast command line utility, with few dependencies, making it straightforward to install and run. It addresses the need for a lightweight tool to rapidly search and extract subsets of reads from raw data and can be easily integrated into comprehensive downstream data analysis pipelines.

## Acknowledgement

We would like to acknowledge Alicia Oshlack for invaluable discussions and feedback on the manuscript. We would also like to thank Givanna Putri, Belinda Phipson and Edward Yang for testing Flexiplex and providing feedback. NMD is funded by NHMRC Investigator Grant GNT2016547.

