## Supplementary for "Flexiplex: A versatile demultiplexer and search tool for omics data"

### Supplementary Methods

#### Flexiplex's Algorithm

Flexiplex is implemented in C++ using Edlib version 1.2.7. A summary of Flexiplex's algorithm is outlined briefly below. The complete source code is open source and available from [https://github.com/DavidsonGroup/flexiplex](https://github.com/DavidsonGroup/flexiplex). All benchmarking was performed with Flexiplex version 1.01.

```
FOR each fastq/fasta entry:
    FOR forward and reverse strands:
        UNTIL no more barcodes identified:
            Search read for the sequence "[left flank][?_b][?_u][right flank]" using
            edlib, where ?_b = a string of '?'s the length of the expected barcode
            and ?_u is a string of '?'s the length of the expected UMI.
            '?' is passed to edlib as a wildcard sequence.
            IF a match is identified within a user specified edit distance
            (-f, default: 8):
                IF barcode discovery mode:
                    Add sequence corresponding to ?_b to the list
                    of identified barcodes
                ELSE:
                    Extract sequence corresponding to ?_b?_u +/- 5bp (target)
                        FOR each known barcode:
                            Calculate the Levenshtein distance
                            between the known barcode and target,
                            penalizing for gaps at the start and end
                            of the known barcode but not the target
                            IF distance is lower than best match:
                                Set as current best match
                            ELSE IF equal to best match:
                                Set barcode as ambiguous
                        IF final best match is within the user defined
                        maximum edit distance (-e, default: 2):
                            Add best matching barcode and
                            corresponding UMI to list of
                            identified barcodes.
            Mask the sequence matching "[left flank][?_b][?_u][right flank]"
    IF barcode demultiplexing mode:
        FOR each identified barcode:
            Print modified read sequence (e.g. adapter trimming,
            read_id supplemented with barcode + UMI depending on user options)
            Print table line for read id, barcode, edit distances and UMI
IF barcode discovery mode:
    Print Barcode frequency tables
```

### Flexiplex-filter's Algorithm

```
Read the list of barcode entries produced by Flexiplex when run in Discovery mode, which
contains each barcode, along with a corresponding number of reads which contain the barcode
(counts)

Sort the list of barcode entries by count

FOR each barcode entry:
       Calculate a rank based on the count, with the entry with the highest count being
       rank 1

FOR each barcode entry, with rank i:
       Calculate an approximate (numerical) derivative for each entry d(rank)/d(count),
       with log-transformed axes:
       [log10(i+1) - log10(i)] ÷ [log10(count[i+1]) - log10(count[i])]
       Calculate an average derivative for each entry using a 'window' of size 9: take the
       mean of the approximate derivatives corresponding to ranks
       [i-4, i-3, i-2, i-1, i, i+1, i+2, i+3, i+4]

Select the barcode entry with the most negative average derivative which:
       - Has a count lower than a user-provided upper bound OR is not ranked in the top 50
       if no upper bound is provided; AND
       - Has a count higher than a user-provided lower bound OR is not ranked below the
       top 95th percentile of entries

FOR each barcode entry with rank better or equal to the selected barcode entry:
       Print the barcode
```

### Fast and accurate sequence search

The single cell sequence search use case used the Illumina short-read single cell RNA-Seq dataset from Chen et al., Cancer Res., 2021 which is available for download from the Sequence Read Archive (SRA) under accession number SRR10971813. Grep (GNU version 2.20), ugrep (version 3.11.2), seqkit grep (version 2.3.0) and Flexiplex (version 1.01), were run on the raw fastq data from the second read pair (R2) using the following commands:

**grep:**

```
grep -h --no-group-separator -F -e $SEQ -e $SEQ_R -B1 -A2
```

**ugrep:**

```
ugrep -Z2 --no-group-separator -F --bool "$SEQ|$SEQ_R" -B1 -A2
```

**seqkit:**

```
seqkit grep -j 1 -m 2 -s -p $SEQ
```

**flexiplex - BCAS4-BCAS3 & EA1:**
```
flexiplex -x $SEQ -d grep -f 2
```

**flexiplex - rs878887783:**
```
flexiplex -f 2 -e 0 -x ${SEQ:0:25} -k ${SEQ:25:5} -x ${SEQ:30:24} -i false
```

Where `SEQ` was one of the following query sequences and `SEQ_R` refers to its reverse complement.

BCAS4-BCAS3 (SNP) - `CCGATCCTGGGGCCGAGGTACCTTTGACAGGAGC`
BCAS4-BCAS3 (reference allele) - `CCGAGCCTGGGGCCGAGGTACCTTTGACAGGAGC`
Adenovirus 5 EA1 - `TTTGGACTTGAGCTGTAAACGCCCCAGGCCATAA`
rs878887783 - `CTCTGATTACTCCTGCCATCATGACCCCTGGCCATAATATGATTTATCTCCACA`

To obtain cellular barcodes we extract the first 16 bases of the matched read pairs, i.e by identifying the read line and ID (`grep "^@"`), searching for the matched read in the R1 read pair file (using `grep -A1`), and extracting the first 16 characters (`cut -c1-16`).

To assess the correspondence of cells with gene expression clustering, a standard Seurat workflow was employed to process the data and visualize the UMAP (Uniform Manifold Approximation and Projection). The raw counts were downloaded from GEO (accession number GSM4285803). A Seurat object was created from the raw counts, followed by the selection of the most variable features (top 2000 features) and data scaling by a linear model. All the cells in the raw counts were retained. We then ran RunPCA (default parameters) and RunUMAP by utilizing 1-20 PCAs to reduce the dimensionality. Subsequently, we ran FindNeighbors (20 PCAs) to find the nearest neighbors of each cell and then FindClusters with a resolution/granularity of 0.2 to identify the cell clusters. Orthogonally, we ran cellSNP-lite (version 1.2.0) on the mapped reads from CellRanger, with a list of common human SNPs provided by the cellSNP-lite authors at https://sourceforge.net/projects/cellsnp/files/SNPlist/. Vireo (version 0.5.8) was then used to cluster the cells into seven donors based on expressed SNPs. Cells that were doublets or unassigned were removed from the analysis. For all remaining cells the SNP-based clustering and gene-expression clustering gave identical groupings. Only one read was required to assign a variant to a cell.

#### Demultiplexing cellular barcode from noisy long-read data

Demultiplexing was assessed using the 'Large' simulation from Ebrahimi et al., iScience 2022, available from [http://10.0.23.196/m9.figshare.19740475.v1](http://10.0.23.196/m9.figshare.19740475.v1), the scmixology 2 dataset available from ENA (accession SAMEA110357667), and the PromethION dataset from You et al., Genome Biology, 2023 available from ENA (accession PRJEB54718).

Corresponding lists of known barcodes were taken from the unique set of true barcodes (L.truth.tsv) and short-read derived barcodes (https://melbourne.figshare.com/articles/dataset/Analysis_data_for_BLAZE/20335203/)

respectively. Flexiplex (version 1.01), scTagger (version 1.1.1) and FLAMES (cloned from https://github.com/LuyiTian/FLAMES github on 7 June 2022) were run on the gzipped fastq files using the following commands, varying the maximum edit distance (ED) between 0 and 3.

**Flexiplex:**
```
gunzip -c $original_fastq_gz | ./flexiplex \
      -d 10x3v3 \
      -e $ED   \
      -k $barcode_file \
      -p $threads \
      | gzip > out.fastq.gz
```

**scTagger:**
```
scTagger.py extract_lr_bc \
      -r $original_fastq_gz \
      -t $threads \
      -z -o temp.out.gz
gunzip temp.out.gz
scTagger.py match_trie \
      -mr $ED \
      -lr temp.out \
      -sr $barcode_file \
      -t $threads \
      -o out.txt
```

**FLAMES:**
```
match_cell_barcode \
       $original_fastq_gz \
       out_stats.txt out.fastq.gz \
       $barcode_file \
       $ED 12
```

For consistency with Flexiplex, which removes ambiguous barcodes (ie. equi-distant best matches), only reads reported by scTagger with a single barcode were compared.

#### Simulation analysis:

To assess the accuracy of each tool on the simulation, we compared the demultiplexed barcodes and their reverse complement against the truth set (L.truth.tsv).

#### Scmixology 2 analysis

For scmixology 2, we used genotyped reads and short-read SNP-clustering to assess barcode correctness. Specifically, we:

1. First constructed a set of cell line specific SNP sequences, by:
    a. Downloading cell line variants from CCLE (https://sites.broadinstitute.org/ccle/datasets)

b. Subsetting to SNPs unique to one of the 5 scmixology cell lines (2479 variants)
c. The SNP +/- 15 bp of sequence from the reference genome (hg19) was extracted using bedtools.
d. These sequences were subsequently searched for using grep on a long read replicate dataset of scmixology (~500 cells) available from SRA (accession SRR12282458), which had been demultiplex prior using Flexiplex and FLAMES.
e. We obtained the cell line annotation of the ~500 cells from prior work (Davidson et al., Genome Biol. 2022).
f. To remove SNPs which were not cell line specific (e.g. due to sequence homology), we filtered for only those with >2 reads and where >90% of cells with the SNP came from the same cell line.
g. SNPs which passed the previous step in both the Flexiplex and FLAMES demultiplexed fastq were kept (92 SNPs). This dataset comes from a different set of cells to scmixology 2, so the choice of demultiplexing tool here should not bias the performance evaluation.

2. Demultiplexed the long read scmixology fastq file using Flexiplex, FLAMES and scTagger for edit distances 0,1,2 and 3, and a range of threads, as described above.
3. SNP-typed the long reads, by searching each read for each of the 92 cell line specific SNPs using grep. For Flexiplex and FLAMES, this was done on the demultipled fastq files. For scTagger, which does not produce a fastq file, we searched the raw fastq files and then mapped read IDs to the barcodes reported by scTagger.
4. For each SNP-typed read we mapped the cellular barcode to a cell line using the cell line annotation derived from short-read data from available at: https://melbourne.figshare.com/articles/dataset/Analysis_data_for_BLAZE/20335203. Doublet cells were removed. We then counted the number of reads for which there was a match/mismatch between the cell line known from the SNP and the cell line derived from the barcode and cell line annotation. The number of reads was then normalized to the number of SNP-typed reads from the raw fastq.

#### PromethION analysis

The PromethION dataset and associated short-read barcodes, published in You et al., Genome Biology, 2023 was downloaded from ENA (accession PRJEB54718). A list of barcodes was constructed by mixing the 1022 short-read barcodes (*true list*) with 10000 barcodes randomly sampled from the 10x Genomic CellRanger whitelist of 3 million possible barcodes (3M-february-2018.txt) (*decoy list*). Flexiplex and scTagger were run on the dataset with maximum threads set to 16. We then counted the number of reads with a reported barcode in the *true list* ($T_{uncorrected}$) or the *decoy list* ($F_{uncorrected}$). As scTagger also reports ambiguous barcodes ie. multiple equi-distant barcodes for a read, we examined its output in two way: i) excluding all reads with ambiguous barcodes from the analysis which is similar to what Flexiplex does and ii) counting ambiguous barcodes towards $T_{uncorrected}$ if any of the reported barcodes were in the *true list*, or $F_{uncorrected}$ if all reported barcodes were in the *decoy list*. To account for false positives where a barcode from the *true list* was reported, we applied the following correction which assumes false positives to the *true list* and *decoy list* are in proportion to their list length:

$F_{corrected} = F_{uncorrected} / r$

$T_{corrected} = T_{uncorrected} - (F_{corrected} - F_{uncorrected})$
Where r is the proportion of barcodes in the *decoy list*.
$T_{correcteded}$ and $F_{corrected}$ were then normalized by the total number of reads in the raw fastq to obtain a final estimate of correct and incorrect demultiplexing rates.

#### Discovering cellular barcodes from noisy long-read data

The four datasets referenced by You et al., Genome Biology, 2023 were used for benchmarking and are available from ENA under accession PRJEB54718. Each of Flexiplex (version 1.01), wf-single-cell (version 1.0.0), BLAZE (version 2.1.4) and scTagger (commit 0b1f9b8) were run on the raw .fastq files, using the commands:

**Flexiplex:**

```
~/flexiplex/flexiplex -f 0 -p $threads $fastq_file > my_barcode_list.txt
```

```
flexiplex-filter --whitelist 3M-february-2018.txt \
                 --outfile filtered_barcodes.txt  \
                 flexiplex_barcodes_counts.txt
```

**BLAZE:**

```
blaze --expect-cells=$expected_cells    \
      --threads=$threads                \
      --no-demultiplexing               \
      $fastq_file
```

**scTagger:**

```
python ~/software/scTagger/scTagger.py extract_lr_bc \
      -t $threads          \
      -r $fastq_file       \
      -o "lr_output.tsv.gz" \
      -p "plots"
```

```
gunzip lr_output.tsv.gz
```

```
python ~/software/scTagger/scTagger.py extract_sr_bc_from_lr \
      -i "lr_output.tsv"       \
      -wl 3M-february-2018.txt \
      -o "sr_output.tsv.gz"
```

```
flexiplex-filter --whitelist 3M-february-2018.txt \
      -u 1000                           \
      --outfile filtered_barcodes.txt   \
      sr_output.tsv
```

**wf-single-cell:**

```
nextflow run epi2me-labs/wf-single-cell  \
      -profile singularity               \
      --fastq $fastq_file                \
      --kit_name 3prime                  \
      --kit_version v3                   \
      --expected_cells $expected_cells   \
      --ref_genome_dir GRCh38-2020-A     \
      --max_threads $threads             \
      -with-timeline
```

The estimated number of cells (`$expected_cells`) passed to BLAZE and wf-single-cell was 200 for scmixology2 and 850 for the three hiPSC datasets. Flexiplex and scTagger both do not require an estimated number of cells, so this information was not provided. However, in order for the inflection point to be correctly identified with flexiplex-filter on hiPSC barcodes from scTagger, the upper search range was set to 1000.

wf-single-cell was run using the Nextflow workflow manager (v23.10.0), using the SLURM executor (v23.02.7) and singularity container (v3.7.4). 20 concurrent executors were allowed and we passed in a maximum thread count of 16 to run the workflow. All of the real time and memory benchmarks for wf-single-cell were obtained from the default timeline generated by Nextflow when the `-with-timeline` parameter is given.

### Computational Benchmarking

Time and memory requirements were recorded with the Linux `/usr/bin/time` command. All use cases were benchmarked on a high performance computing cluster (Intel® Xeon® CPU E5-2690 v4 @ 2.60GHz) running CentOS 7. Although Flexiplex allows multiple threads, we benchmarked demultiplexing and search performance using a single thread unless stated otherwise. This allowed a fair evaluation against the non-threaded methods, grep and FLAMES (match_cell_barcode). For cellular barcode discovery, we benchmarked each dataset with 1, 2, 4, 8, and 16 threads, to compare performance across Flexiplex, scTagger, and BLAZE.

### Supplementary Tables

| **Cell Line** | **Tool** | **BCAS4-BCAS3 (SNP)** MCF7 | **BCAS4-BCAS3 (Ref.)** MCF7 | **rs878887783** T47D | **Adenovirus** HEK293T |
|---|---|---|---|---|---|
| MCF7<br>177 million reads | Flexiplex | **4356** | **4123** | 6 | 0 |
| | Grep | 3599 | 4 | **1** | 0 |
| | Ugrep | 4273 | 4064 | 7419 | 0 |
| | Seqkit grep | 4175 | 4030 | 7378 | 0 |
| T47D<br>103 million reads | Flexiplex | 0 | 0 | 26 | 0 |
| | Grep | 0 | 0 | 8 | 0 |
| | Ugrep | 0 | 0 | **27** | 0 |
| | Seqkit grep | 0 | 0 | 21 | 0 |
| MDAMB134 VI<br>126 million reads | Flexiplex | 0 | 0 | **0** | 0 |
| | Grep | 0 | 0 | **0** | 0 |
| | Ugrep | 0 | 0 | 2872 | 0 |
| | Seqkit grep | 0 | 0 | 2879 | 0 |

***Supplementary Table 1:*** *To confirm that false positives seen in single-cell data were unrelated to errors in sequence searching, we also examine matched bulk RNA-Seq cell line data. Three of the seven cell lines were available from the Cancer Cell Line Encyclopedia (CCLE) (SRA Accessions: SRR8615755, SRR8615758 and SRR8615812). Similar to our single-cell analysis, Flexiplex, ugrep and seqkit were run allowing an edit distance up to two, whereas grep required a perfect match. Cell lines for which a variant is expected to be found are highlighted in green. Flexiplex identified the highest or second highest number of reads for both variants. For BCAS4-BCAS3 and the Adenoviral sequence, no false positives (variants identified in other cell lines) were found. False positives for rs878887783 were found in all cell lines and with all tools. rs878887783 is a point mutation and therefore has lower specificity than BCAS4-BCAS3 or Adenoviral sequence. Flexiplex reported the second lowest number of false positives amongst the tools tested. The high number of false positives reported by ugrep and seqkit are due to detection of the reference allele.*

| Dataset | Tool | Parameters | User time | System time | Mem (GB) | Real time |
|---|---|---|---|---|---|---|
| scmixology2 | Flexiplex | 1 thread | 0:15:21 | 0:00:15 | 1.47 | 0:15:37 |
| | | 2 threads | 0:15:29 | 0:00:18 | 1.47 | 0:08:40 |
| | | 4 threads | 0:15:32 | 0:00:16 | 1.46 | 0:05:06 |
| | | 8 threads | 0:15:31 | 0:00:30 | 1.46 | 0:03:27 |
| | | 16 threads | 0:18:47 | 0:00:34 | 1.47 | 0:03:18 |
| | BLAZE | 1 thread | 3:37:25 | 0:00:38 | 1.14 | 3:39:32 |
| | | 2 threads | 3:39:28 | 0:00:36 | 1.14 | 3:41:39 |
| | | 4 threads | 3:36:44 | 0:01:11 | 1.13 | 1:14:08 |
| | | 8 threads | 3:45:05 | 0:01:17 | 1.14 | 0:34:04 |
| | | 16 threads | 6:20:35 | 0:01:24 | 1.16 | 0:28:44 |
| | sctagger | 1 thread | 0:46:53 | 0:01:43 | 39.39 | 0:46:38 |
| | | 2 threads | 0:47:17 | 0:02:00 | 39.39 | 0:29:01 |
| | | 4 threads | 0:47:47 | 0:02:28 | 39.39 | 0:20:24 |
| | | 8 threads | 0:49:19 | 0:03:18 | 39.39 | 0:16:22 |
| | | 16 threads | 1:07:32 | 0:06:17 | 39.39 | 0:16:00 |
| | wf-single-cell | up to call_adapter_scan | - | - | 37.96 | 1:15:09 |
| | | up to extract_barcodes | - | - | 21.74 | 9:58:46 |

| Dataset | Tool | Parameters | User time | System time | Mem (GB) | Real time |
|---|---|---|---|---|---|---|
| gridION | Flexiplex | 1 thread | 0:05:11 | 0:00:05 | 1.42 | 0:05:16 |
| | | 2 threads | 0:05:21 | 0:00:06 | 1.42 | 0:02:57 |
| | | 4 threads | 0:05:22 | 0:00:09 | 1.42 | 0:01:47 |
| | | 8 threads | 0:05:32 | 0:00:07 | 1.42 | 0:01:15 |
| | | 16 threads | 0:07:14 | 0:00:12 | 1.42 | 0:01:06 |
| | BLAZE | 1 thread | 0:57:08 | 0:00:11 | 1.18 | 0:57:51 |
| | | 2 threads | 0:58:10 | 0:00:11 | 1.18 | 0:58:36 |
| | | 4 threads | 1:01:17 | 0:00:25 | 1.19 | 0:22:12 |
| | | 8 threads | 1:03:34 | 0:00:33 | 1.19 | 0:13:13 |
| | | 16 threads | 1:31:14 | 0:00:36 | 1.20 | 0:09:31 |
| | sctagger | 1 thread | 0:13:08 | 0:00:43 | 13.16 | 0:13:03 |
| | | 2 threads | 0:13:12 | 0:00:48 | 13.16 | 0:08:31 |
| | | 4 threads | 0:13:25 | 0:00:57 | 13.16 | 0:06:21 |
| | | 8 threads | 0:13:40 | 0:01:12 | 13.16 | 0:05:22 |
| | | 16 threads | 0:18:00 | 0:01:58 | 13.16 | 0:05:13 |
| | wf-single-cell | up to call_adapter_scan | - | - | 16.40 | 0:27:57 |
| | | up to extract_barcodes | - | - | 19.45 | 3:43:48 |

| Dataset | Tool | Parameters | User time | System time | Mem (GB) | Real time |
|---|---|---|---|---|---|---|
| gridION-q20 | Flexiplex | 1 thread | 0:03:36 | 0:00:05 | 1.39 | 0:03:40 |
| | | 2 threads | 0:03:37 | 0:00:05 | 1.39 | 0:02:03 |
| | | 4 threads | 0:03:38 | 0:00:05 | 1.39 | 0:01:15 |
| | | 8 threads | 0:03:42 | 0:00:05 | 1.39 | 0:00:53 |
| | | 16 threads | 0:04:59 | 0:00:08 | 1.39 | 0:00:53 |
| | BLAZE | 1 thread | 0:23:53 | 0:00:06 | 1.16 | 0:24:09 |
| | | 2 threads | 0:24:55 | 0:00:06 | 1.16 | 0:25:16 |
| | | 4 threads | 0:26:03 | 0:00:14 | 1.19 | 0:09:30 |
| | | 8 threads | 0:27:03 | 0:00:14 | 1.19 | 0:05:30 |
| | | 16 threads | 0:39:38 | 0:00:20 | 1.19 | 0:04:09 |
| | sctagger | 1 thread | 0:06:00 | 0:00:23 | 5.98 | 0:05:57 |
| | | 2 threads | 0:06:09 | 0:00:26 | 5.99 | 0:04:08 |
| | | 4 threads | 0:06:11 | 0:00:31 | 5.99 | 0:03:12 |
| | | 8 threads | 0:06:18 | 0:00:36 | 5.99 | 0:02:45 |
| | | 16 threads | 0:08:00 | 0:01:01 | 5.99 | 0:02:43 |
| | wf-single-cell | up to call_adapter_scan | - | - | 15.74 | 0:13:29 |
| | | up to extract_barcodes | - | - | 19.17 | 1:49:31 |

| Dataset | Tool | Parameters | User time | System time | Mem (GB) | Real time |
|---|---|---|---|---|---|---|
| promethION | Flexiplex | 1 thread | 0:50:11 | 0:00:48 | 1.61 | 0:50:58 |
| promethION | Flexiplex | 2 threads | 0:54:36 | 0:00:49 | 1.61 | 0:31:31 |
| promethION | Flexiplex | 4 threads | 1:02:17 | 0:00:48 | 1.61 | 0:26:06 |
| promethION | Flexiplex | 8 threads | 1:02:06 | 0:00:51 | 1.61 | 0:19:21 |
| promethION | Flexiplex | 16 threads | 1:08:03 | 0:02:18 | 1.61 | 0:12:52 |
| promethION | BLAZE | 1 thread | 7:42:41 | 0:01:17 | 1.40 | 7:47:25 |
| promethION | BLAZE | 2 threads | 7:39:35 | 0:01:12 | 1.40 | 7:43:25 |
| promethION | BLAZE | 4 threads | 7:55:20 | 0:02:53 | 1.37 | 2:46:53 |
| promethION | BLAZE | 8 threads | 8:42:21 | 0:04:20 | 1.38 | 1:45:28 |
| promethION | BLAZE | 16 threads | 12:47:36 | 0:05:01 | 1.41 | 1:17:35 |
| promethION | sctagger | 1 thread | 2:40:47 | 0:04:54 | 103.95 | 2:41:28 |
| promethION | sctagger | 2 threads | 2:41:46 | 0:06:09 | 103.95 | 1:35:37 |
| promethION | sctagger | 4 threads | 2:44:36 | 0:08:07 | 103.95 | 1:03:51 |
| promethION | sctagger | 8 threads | 2:57:46 | 0:11:33 | 104.90 | 0:48:43 |
| promethION | sctagger | 16 threads | 4:21:59 | 0:21:06 | 103.95 | 0:47:33 |
| promethION | wf-single-cell | up to call_adapter_scan | - | - | 81.16 | 3:36:26 |
| promethION | wf-single-cell | up to extract_barcodes | - | - | 20.31 | 28:50:46 |

***Supplementary Table 2:*** *Computational performance of Flexiplex, BLAZE, sctagger, and wf-single-cell for barcode discovery. Flexiplex consistently ran the fastest, with only marginally higher memory requirements than BLAZE. The scTagger benchmark does not include the time taken to run flexiplex-filter; in practice, flexiplex-filter had minimal on total runtime, and took under 10 seconds on all the datasets tested. The wf-single-cell pipeline requires full alignment with the reference human genome in order to produce a barcode list, however, none of the other tools tested required an alignment. This time is indicated by "up to extract_barcodes". A more comparable metric for computational efficiency is the "up to call_adapter_scan" function, which records only until the read adapter and TSO sequences are located and oriented. This is more comparable to the computations performed by Flexiplex, BLAZE, and scTagger.*

### Supplementary Figures

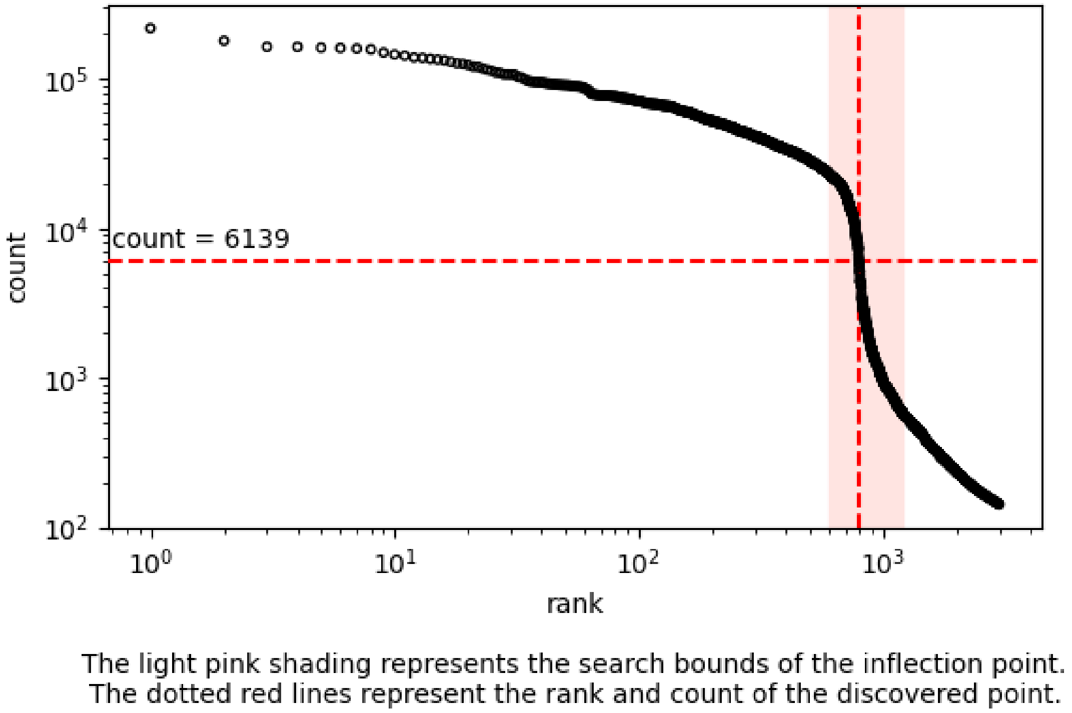

***Supplementary Figure 1:*** *Example knee plot generated by Flexiplex-filter showing the barcode read count (y-axis) against ordered rank (x-axis). The inflection point of the distribution (red dashed lines) can be used to identify the rank below which barcodes should be filtered out. Finding the inflection point and filtering is performed automatically by Flexiplex-filter. This example was made using a barcode list generated by scTagger on the PromethION dataset from* You et al., Genome Biology, 2023.

#### A) BCAS4-BCAS3

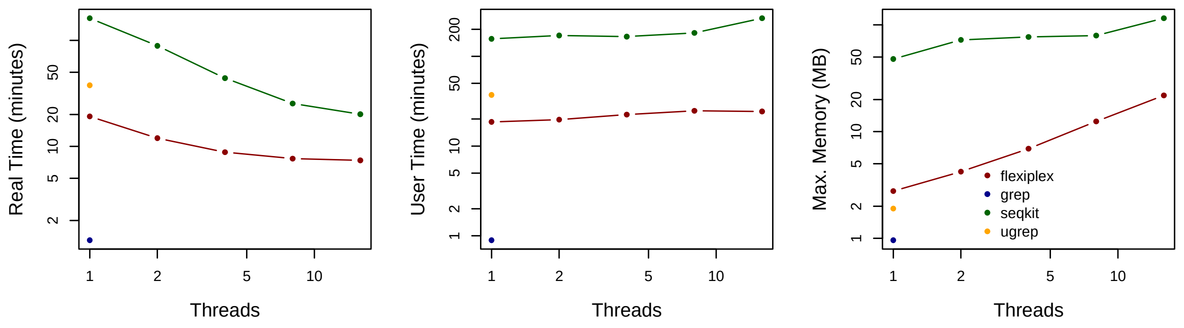

## B) rs878887783

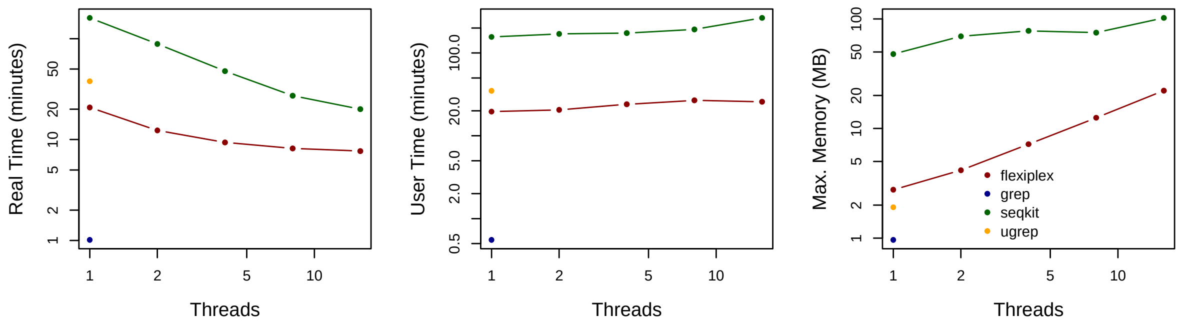

***Supplementary Figure 2:*** *Computational performance of four tools searching for sequence from (A) the BCAS4-BCAS3 fusion and (B) rs878887783. Computational performance for ADENO virus is not shown as it is similar to BCAS4-BCAS3. Grep and ugrep could only be run with a single thread. Seqkit and Flexiplex were run with 1, 2, 4, 8 and 16 threads.*

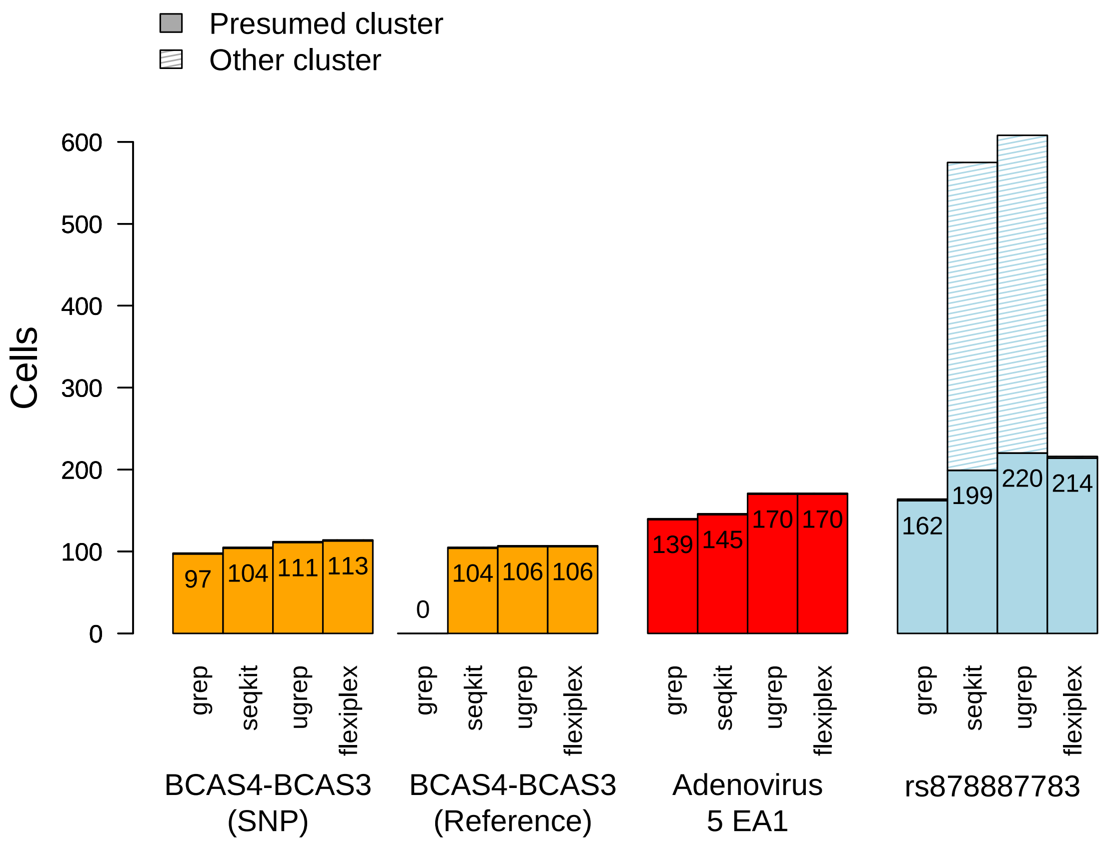

***Supplementary Figure 3:*** *The impact of requiring >1 variant UMI per cell. The false positive rate (`Other cluster` hits) are reduced from up to 9% (Adenovirus) to below 1% for Flexiplex, however the number of cells which can be correctly genotyped is reduced from 47% to 18%. The false positive rate for rs878887783 remains high for seqkit and ugrep as these are caused by detection of the reference allele sequence.*

#### A) Simulation

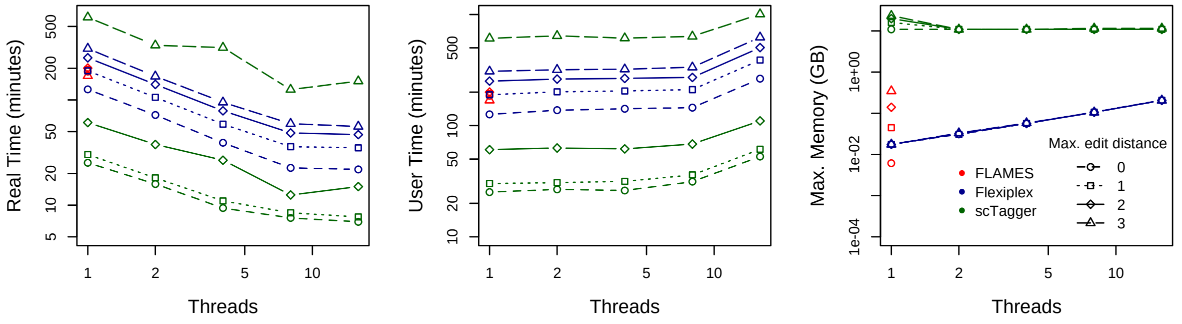

#### B) Scmixology 2

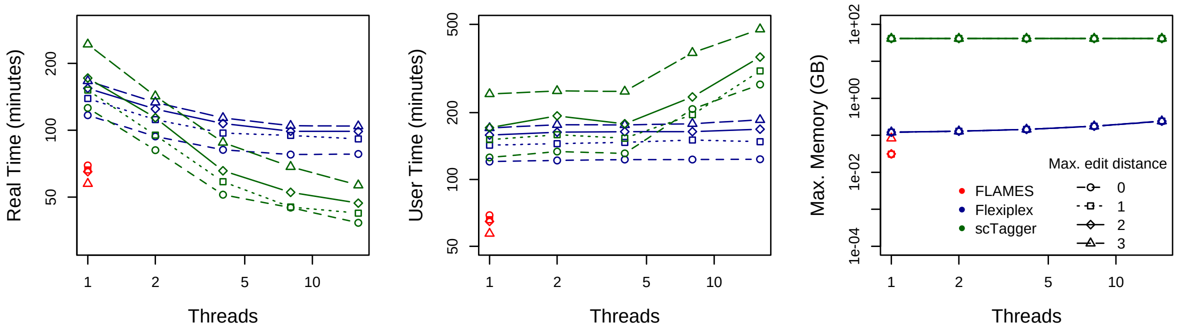

***Supplementary Figure 4***: *Computational performance of three tools for cellular barcode demultiplexing on (A) the simulation dataset and (B) the scmixology 2 dataset. FLAMES could only be run with a single thread. scTagger and Flexiplex were run with 1, 2, 4, 8, and 16 threads. We tested all tools over a range of maximum edit distances, 0t to 3.*
